# An homeotic post-transcriptional network controlled by the RNA-binding protein RBMX

**DOI:** 10.1101/400366

**Authors:** Paola Zuccotti, Daniele Peroni, Valentina Potrich, Alessandro Quattrone, Erik Dassi

## Abstract

Post-transcriptional regulation (PTR) of gene expression is a powerful determinant of protein levels and cellular phenotypes. The 5’ and 3’ untranslated regions of the mRNA (UTRs) mediate this role through sequence and secondary structure elements bound by RNA-binding proteins (RBPs) and noncoding RNAs. While functional regions in the 3’UTRs have been extensively studied, the 5’UTRs are still relatively uncharacterized. To fill this gap, here we used a computational approach based on phylogenetic conservation to identify hyper-conserved elements in human 5’UTRs (5’HCEs). Our assumption, supported by the recovery of functionally characterized elements, was that 5’HCEs would represent evolutionarily stable and hence important PTR sites.

We identified over 5000 short, clustered 5’HCEs occurring in approximately 10% of human protein-coding genes. Among these, homeotic genes were highly enriched. Indeed, 52 of the 258 characterized homeotic genes contained at least one 5’HCE, including members of all four Hox clusters and several other families. Homeotic genes are essential transcriptional regulators. They drive body plan and neuromuscular development, and the role of PTR in their expression is mostly unknown. By integrating computational and experimental approaches we then identified the RBMX RNA-binding protein as the initiator of a post-transcriptional cascade regulating many such homeotic genes. RBMX is known to control its targets by modulating transcript abundance and alternative splicing. Adding to that, we observed translational control as a novel mode of regulation by this RBP.

This work thus establishes RBMX as a versatile master controller of homeotic genes and of the developmental processes they drive.

## Introduction

Post-transcriptional control of gene expression (PTR) has recently emerged as a key determinant of protein levels and the consequent cell phenotypes (Schwanhäusser et al. 2011; Vogel et al. 2010). The two untranslated regions of the mRNA, the 5’ and 3’UTR, mediate this role through sequence and secondary structure elements. These are bound by trans-factors such as RNA-binding proteins (RBPs) and noncoding RNAs (ncRNAs). These factors ultimately control the fate of a transcript by regulating its stability, localization, translation and influencing several other aspects of PTR (Glisovic et al. 2008; Noh et al. 2018; Bartel 2018). RBPs, in particular, are a major player in PTR, counting over 1500 human genes (Gerstberger et al. 2014). Their combined action forms a complex regulatory network of cooperative and competitive interactions (Dassi 2017).

Many works have focused on characterizing regulatory elements in 3’UTRs, mainly concerning mRNA stability and localization. However, less is known about functional regions in 5’UTRs. Which are the factors binding them, and which is their impact on the fate of the transcripts bearing them? A comprehensive catalog of functional regions in the 5’UTRs is still missing, thus hampering our ability to understand the mechanisms exploiting this regulatory hotspot. We and others have successfully compiled such a catalog in 3’UTRs, by exploiting the intuitive concept that evolutionarily conserved regions in mRNAs may be functional (Dassi et al. 2013; Bejerano 2004; McCormack et al. 2012; Reneker et al. 2012; Sathirapongsasuti et al. 2011). The 5’ and 3’UTRs appear to have different functions (Mayr 2017; Hinnebusch et al. 2016), and are thus likely endowed with distinct profiles of cis-elements and targeting trans-factors. However, this phylogenetic approach is general.

Homeotic genes are a key class of developmental regulators (Philippidou and Dasen 2013; McGinnis and Krumlauf 1992; Olson and Rosenthal 1994). They are extremely conserved throughout evolution and found in a range of species going from fungi to mammals (Holland 2013). These proteins, acting as transcription factors, are responsible for defining embryonic regions identity along the anteroposterior and limb axes of vertebrates (Mallo et al. 2010; Gehring 2012). Furthermore, homeotic genes have also been implicated in organ, neural and muscular development (Philippidou and Dasen 2013; Zagozewski et al. 2014; Cambier et al. 2014). Much is known about how their expression is regulated at the transcriptional level (Mallo and Alonso 2013). A few works have also focused on finding regulatory elements in homeotic 3’UTRs, uncovering several mechanisms mediated by RBPs (Nie et al. 2015; Fritz and Stefanovic 2007; Pereira et al. 2013; Rogulja-Ortmann et al. 2014) and miRNAs (Yekta et al. 2004; Li et al. 2014; Wang et al. 2014). Concerning their 5’UTR, however, there is limited evidence about the use of alternative transcription initiation sites (Regadas et al. 2013) and regulation by RNA-binding proteins (Nie et al. 2015). Additionally, IRES-like elements found in the 5’UTR of some Hox genes were observed to control ribosome specificity by recruiting *RPL38 (Xue et al. 2015)*. Globally, we still know little about the homeotic genes 5’UTRs and the post-transcriptional mechanisms acting on them.

Among the potential post-transcriptional regulators of these genes, *RBMX* is a scarcely studied RBP whose inactivation has been previously associated with neuromuscular developmental defects in *X. laevis* (Dichmann et al. 2008) *and D.rerio* (Tsend-Ayush et al. 2005)*. This suggests it as a promising candidate regulator of homeotic genes. RBMX*, also known as *hnRNP G*, belongs to the *RBMY* gene family, of which it is an X-chromosome homolog. It contains a single RRM domain and a C-terminal low-complexity region, by which it binds RNA. It was observed to regulate splice site selection (Heinrich et al. 2009; Wang et al. 2011) and upregulate the expression of the tumor suppressor *TXNIP* by an unspecified mechanism (Shin et al. 2008). Further work found *RBMX* to be involved with the DNA-damage response (Adamson et al. 2012) and regulate centromere biogenesis (Cho et al. 2018; Matsunaga et al. 2012). Recently, N6-methyladenosine marks were found to increase the accessibility of *RBMX* binding sites, thus mediating its effect on its targets expression and splicing (Liu et al. 2017). Eventually, RBMX mutations were associated with the insurgence of an X-linked intellectual disability (XLID) syndrome associated with craniofacial dysmorphisms (Shashi et al. 2015).

We present here the analysis of hyper-conserved regions (HCEs) in 5’UTRs, based on a broad set of 44 vertebrates. Among the 5248 identified HCEs are several known and highly conserved regulatory sites, such as the iron response element (IRE), thus confirming the reliability of our approach. Through this potentially functional catalog of 5’UTR regions, we identified a group of homeotic genes controlled by the *RBMX* RNA-binding protein. We thus describe a novel regulatory network controlling homeotic genes at the post-transcriptional level, and a novel role for *RBMX* as a master translational controller of development.

## Results

### 5’UTR HCEs are short and clustered phylogenetic footprints

To extract phylogenetically conserved regions from the 5’UTR of human mRNAs, we applied to the 5’UTRs the pipeline we previously described for 3’UTRs (Dassi et al. 2013). Briefly, we computed a per-nucleotide hyper-conservation score (HCS), ranging from 0 to 1. The HCS is the average of sequence conservation and the fraction of the phylogenetic tree covered by that UTR alignment (branch length score). A sliding-window approach was then used to find 5-nucleotides seeds with maximum conservation. Seeds were eventually extended upstream and downstream along the UTR while HCS held above a threshold of 0.85, thus extracting only highly conserved regions (see Methods). We term these regions 5’ hyper-conserved elements (5’HCEs).

This approach led to the identification of 5248 HCEs (**Supplementary Table 1**), contained in the 5’UTRs of 2737 transcripts coding for 2228 distinct genes. As shown in **Figure 1A**, these HCEs are mostly short, with an average length of 72 nucleotides and a median of 24 (minimum length 5 nucleotides, maximum 1542). 5’HCEs are thus 28% shorter on average than 3’UTR HCEs (Dassi et al. 2013). However, one should consider that 5’UTRs are also on average almost three times shorter than 3’UTRs (mean length of 455 nts for 5’UTR and 1282 nts for 3’UTRs). We then analyzed the nucleotide composition of 5’HCEs. **Figure 1B** shows the absence of imbalance in the frequencies of AU (typically enriched in 3’UTRs) and GC nucleotides. To further characterize the properties of 5’HCEs, we also observed their relative positional distribution. As can be seen in **Figure 1C**, almost a third of all 5’HCEs (1463/5248) are within 10 bases of another 5’HCE. This figure increases to almost 50% if we exclude 2028 isolated 5’HCEs (i.e., only 5’HCE found on that given 5’UTR), and only a few (155) are at least 50 nucleotides away from another 5’HCE. Globally, if considering a maximum distance of 20 nucleotides between 5’HCEs, 2672 out of 3220 non-isolated 5’HCEs (82%) are found in clusters. This suggests the prevalence of a clustered 5’HCE organization, a pattern also observed for 3’UTR HCEs (Dassi et al. 2013). Eventually, we analyzed the position of 5’HCEs in the containing UTRs. We observed them to be spread along the 5’UTR with a preference for its initial 10% (**Figure 1D**, 48% of HCEs). However, 27% of the 5’HCEs cover 95% or more of their 5’UTR, and thus start around its first bases. When excluding these, this positional preference decreases to 18% of the 5’HCEs only.

**Figure 1:**
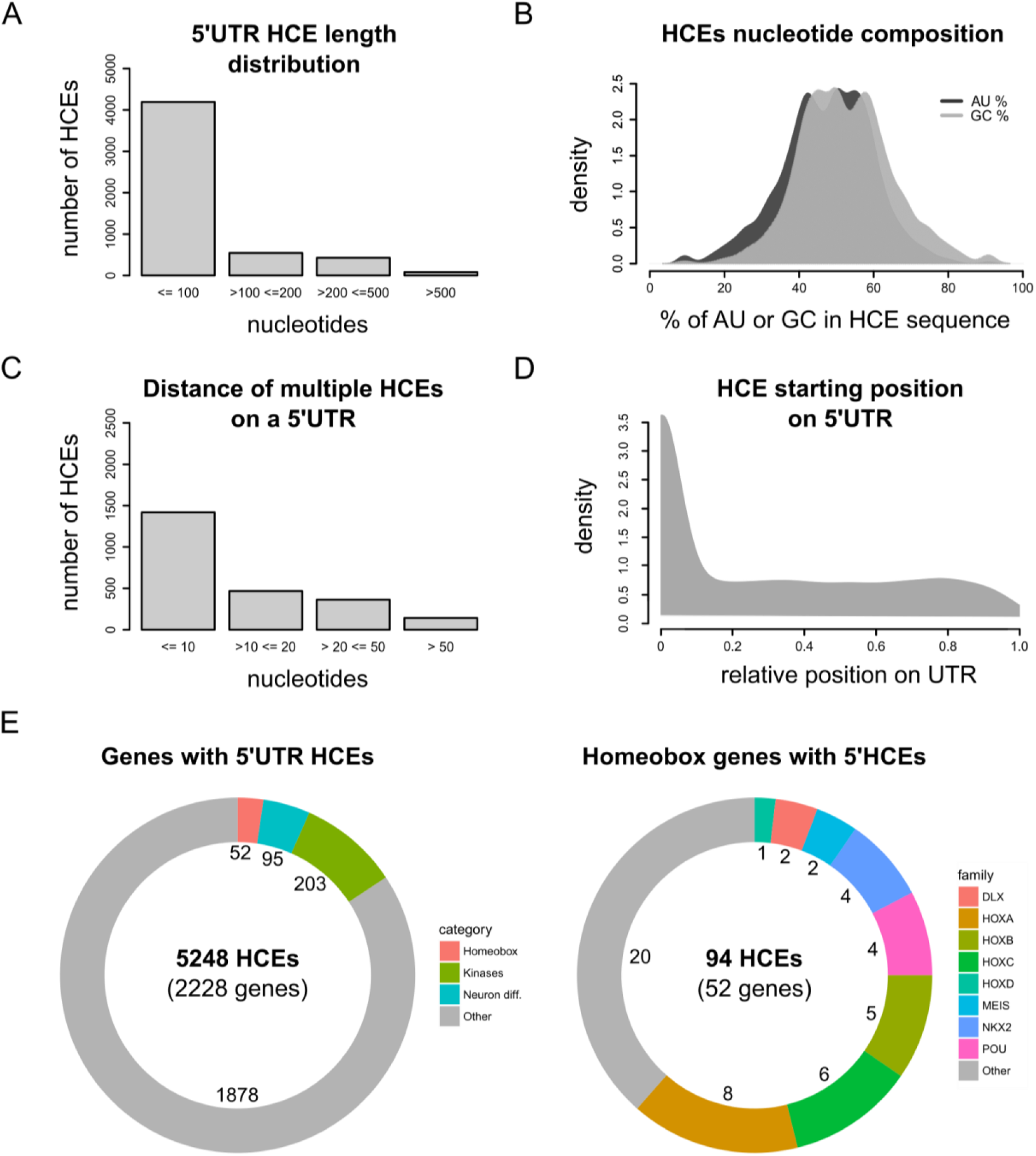
Homeotic genes are enriched in 5’UTR HCEs. **A)** shows the distribution of 5’ hyper-conserved elements (5’HCEs) lengths, highlighting a striking majority of these to be shorter than 100 nucleotides. **B)** displays the density of AU and GC nucleotides frequencies in 5’HCEs, indicating that no significant bias in the composition can be observed. **C)** shows the distance between HCEs on 5’UTRs carrying more than one of these elements. There is a tendency for HCEs to be “clustered”, i.e. in proximity to one another. **D)** displays the density of relative HCE start positions on 5’UTRs (0 = UTR start, 1 = UTR end). HCEs are evenly distributed with the exception of an increased density in the first 10% of the UTR (representing less than 25% of all HCEs). **E)** shows the distribution of 5’HCEs by functional gene categories (left) and the abundance of homeotic families in HCE-containing homeobox genes (right). Numbers next to each category/family show the related number of genes.

We then sought to understand whether the identified 5’HCEs are representative of functional regions in the 5’UTRs. We thus searched in the AURA2 database (Dassi et al. 2014) for several 5’UTR cis-elements and RBP binding sites that were previously characterized and are known to be highly conserved. In particular, as shown in **Supplementary Figure 1**, we first considered two conserved iron response elements (IREs) (Gray et al. 1996; Hentze et al. 1987) in *ferritin* (*FTL*, **S1A**) and *aconitase 2* (*ACO2*, **S1B**) mRNAs. In both cases, an HCE is identified in that 5’UTR and contains the IRE. We then considered two conserved binding sites for the *LARP6* RBP, on *collagen alpha type 1* (*COL1A1*, **S1C**) (Cai et al. 2010) and *ornithine decarboxylase 1* (*ODC1*, **S1D**) (Manzella and Blackshear 1992) mRNA. While for *COL1A1* the identified HCE completely contains the *LARP6* binding site, in the case of *ODC1* the overlap is only partial but still present. Furthermore, 5’HCEs do not overlap with uORFs (Wethmar 2014) or IRESs (Yamamoto et al. 2017). Globally, these observations show that our HCE detection algorithm can identify functionally relevant phylogenetic footprints in 5’UTRs. Furthermore, it suggests 5’HCEs to be potential binding sites for RBPs.

### Homeotic genes are enriched in 5’HCEs

We then annotated groups of functionally related genes among the 2228 containing one or more 5’HCEs. To do so, we performed a functional enrichment of genes, pathways and protein domain ontologies using DAVID (Huang et al. 2007). The results we obtained, as shown in **Figure 1E** and **Supplementary Table 2**, revealed the presence of three functional themes endowed with high significance. The first theme, the homeobox, involves 52 genes representing several families of these essential transcription factors (Ladam and Sagerström 2014). Homeobox genes are responsible for developmental patterns (Gehring 2012; Philippidou and Dasen 2013; Zagozewski et al. 2014) and are highly conserved throughout vertebrates, from the fruit fly to human (Holland 2013). A second, functionally broader theme, regroups 95 genes implicated in neuronal differentiation, some of which are also part of the previous theme. The last theme is made up of 203 protein kinases. These include various kinase types (Ser/Thr, Tyr, and others) affecting several signaling pathways (such as MAPK, NFKB, and others, as shown in **Figure 1E**). Given the importance of homeotic genes in development and their high functional coherence, we decided to focus our attention on this theme. We reasoned that these features could allow us to trace a meaningful homeobox PTR network and investigate the related regulatory mechanisms.

Among these 52 genes, whose 5’UTRs contain 94 HCEs, we can find members of all four Hox clusters (eight *HOXA*, five *HOXB*, six *HOXC* and one *HOXD* genes). Other families are also included, such as *NKX* and *POU* (four genes each), *MEIS* and *DLX* (two genes each). All these proteins contain a homeobox domain and control developmental processes. Nevertheless, specific functions such as pattern specification (25 genes), cell motion (10 genes) and neuronal differentiation (20 genes) involve only a subset of these 52 genes. Homeobox 5’HCEs have a median length of 28 nucleotides (ranging from 5 to 423 nucleotides). They are often clustered, as 48/63 non-isolated HCEs are within 20 nucleotides of one another. **Supplementary Table 3** and **4** present the complete list of genes composing this theme and their functional annotation.

### Homeotic 5’HCEs contain an RBMX-binding signature

We thus asked ourselves whether this set of homeotic genes could be controlled by a common regulatory mechanism through binding sites within their 5’HCEs. To identify such mechanism we first performed an integrative motif search with DynaMIT (Dassi and Quattrone 2016), combining a sequence search (using Weeder (Pavesi et al. 2004)) with an RNA secondary structure search (using RNAforester (Höchsmann et al. 2003)). Integration of the motifs identified by the two tools was done by clustering motifs co-occurring on the same sequences. Among the results, the best cluster included both sequence and secondary structure motifs shared by most homeotic 5’HCEs. The resulting motif, as shown in **Figure 2A**, is short, unstructured, and C-rich. Breaking down the consensus by its composing motifs reveals CGAC as shared by sequence search motifs of all length and CCAG as secondary structure search consensus.

**Figure 2:**
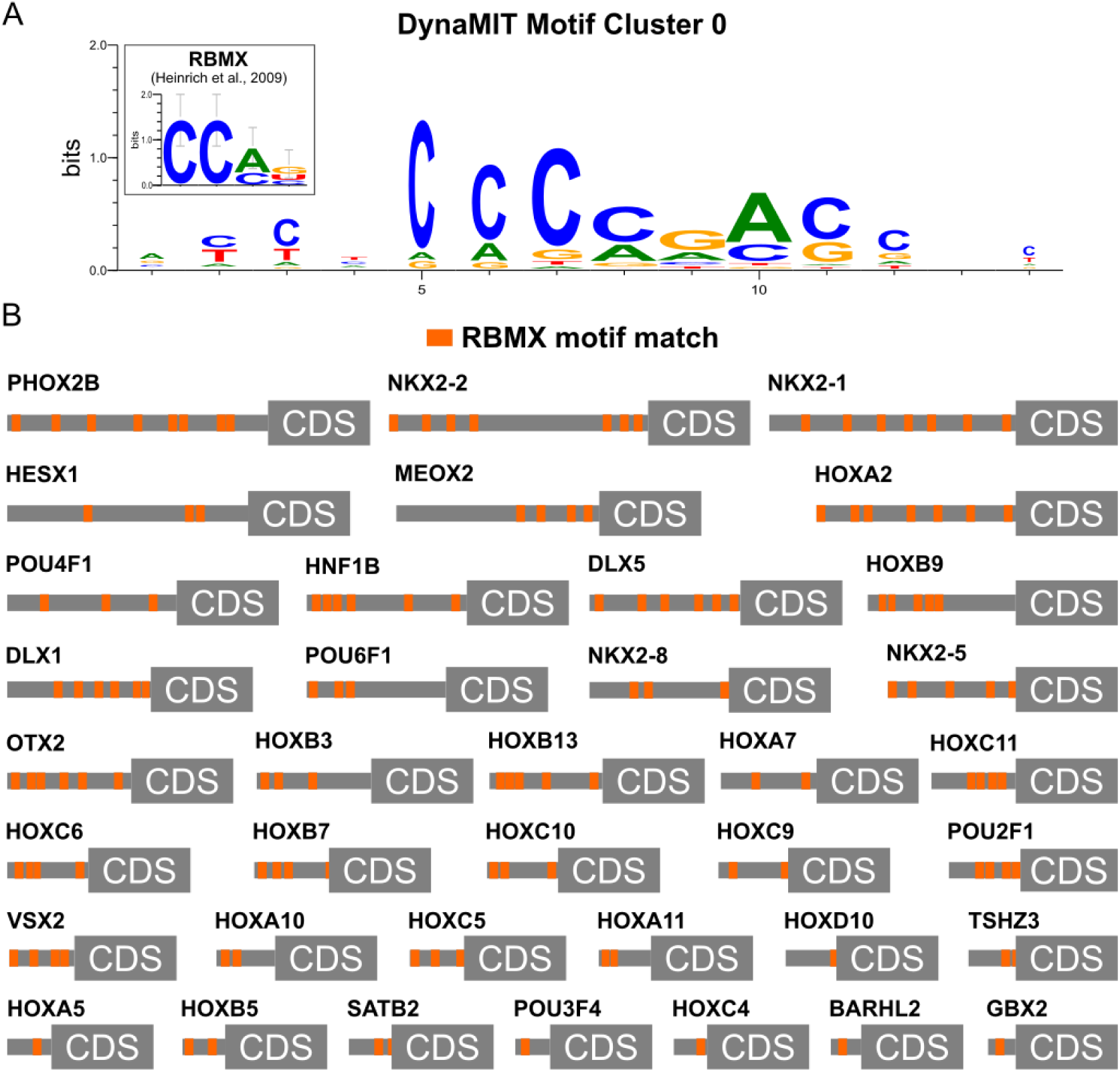
Homeotic genes 5’HCEs contain an RBMX binding signature. **A)** presents the best motif cluster identified by DynaMIT in homeotic 5’HCEs by integrating sequence and secondary structure motif searches. The leftmost inset displays the currently known binding motif of RBMX as a WebLogo for comparison. **B)** displays matches for the RBMX binding motif on the portion of homeotic genes 5’UTRs contained into an HCE. Matches, represented by orange boxes, are clustered in 15 nucleotides windows (i.e. a single orange box may include multiple matches within 15 nucleotides) for visualization purposes.

Given this motif indication, we then proceeded by trying to understand which trans-factor may be binding it in order to exert a regulatory function on these homeotic genes. To this goal, we performed a search on known RBP binding motifs using the CISBP-RNA database (Ray et al. 2013). The results highlighted a protein, RBMX, having a binding consensus strikingly similar to the motifs found in these 5’HCEs (similarity score 93.5% and Pearson correlation 0.73). Its known consensus, derived by a SELEX experiment (Heinrich et al. 2009) is shown as a weblogo in the inset of **Figure 2A**.

To systematically map potential RBMX binding sites on homeotic genes 5’HCEs, we thus performed a pattern match analysis with the RBMX binding motif. The results, displayed in **Figure 2B**, show potential RBMX binding sites distributed throughout the homeotic genes 5’UTRs. These sites appear to be preferentially located in the proximity of each other (median 17 nts, average 41). Such sites distribution hints to a potential for homomultimeric RNA binding as previously observed for RBMX (Heinrich et al. 2009).

### RBMX binds to homeotic genes mRNAs

The motif signature we detected in 5’HCEs of homeotic genes suggests that RBMX may contribute to the post-transcriptional regulation of their mRNAs. *RBMX* (also known as *HNRNPG*) is an RBP of the *RBMY* family, associated with neuromuscular developmental defects in *X. laevis* (Dichmann et al. 2008) and *D. rerio* (Tsend-Ayush et al. 2005), thus making it a promising candidate for post-transcriptional control of homeotic genes. To investigate this assumption, we first sought to confirm that RBMX is binding to homeotic mRNAs. We performed an RNA immunoprecipitation (RIP) assay followed by targeted RNA-sequencing in HEK293 cells, thus probing RBMX binding strength on the mRNAs of 50 genes. These included our 38 “core” homeotic genes, RBMX, two controls and additional members of families represented in the core genes. We computed the fold enrichment for each gene as the ratio between the normalized abundance in the RIP and the corresponding input samples. As shown in **Figure 3A**, we found 29/38 (76%) core homeotic genes to be enriched at least twofold in at least two of the three replicates (23/38 if using a fold enrichment threshold of four). Members of all four HOX clusters and other families such as *NKX* and *POU* were enriched. If considering all the 50 tested genes, 40 (80%) are enriched at least twofold in at least two replicates (32/50 if using a fold enrichment threshold of four). Among these, we find a known target of RBMX (Heinrich et al. 2009), FYN, which we used as positive control (2/3 replicates, average fold enrichment of 5.45). Our negative control, HNRNPM, is instead not enriched in any replicate (average fold enrichment of 1.43), as shown in **Figure 3B**. Remarkably, RBMX also binds to its cognate mRNA (all replicates, median fold enrichment 8.11), a frequent feature of RBPs (Dassi 2017). The list of bound genes can be found in **Supplementary Table 5**. Globally, this assay suggests that RBMX is indeed binding to a wide set of homeotic genes mRNAs. Thus, this RBP likely contributes to the post-transcriptional regulation of their expression.

**Figure 3.**
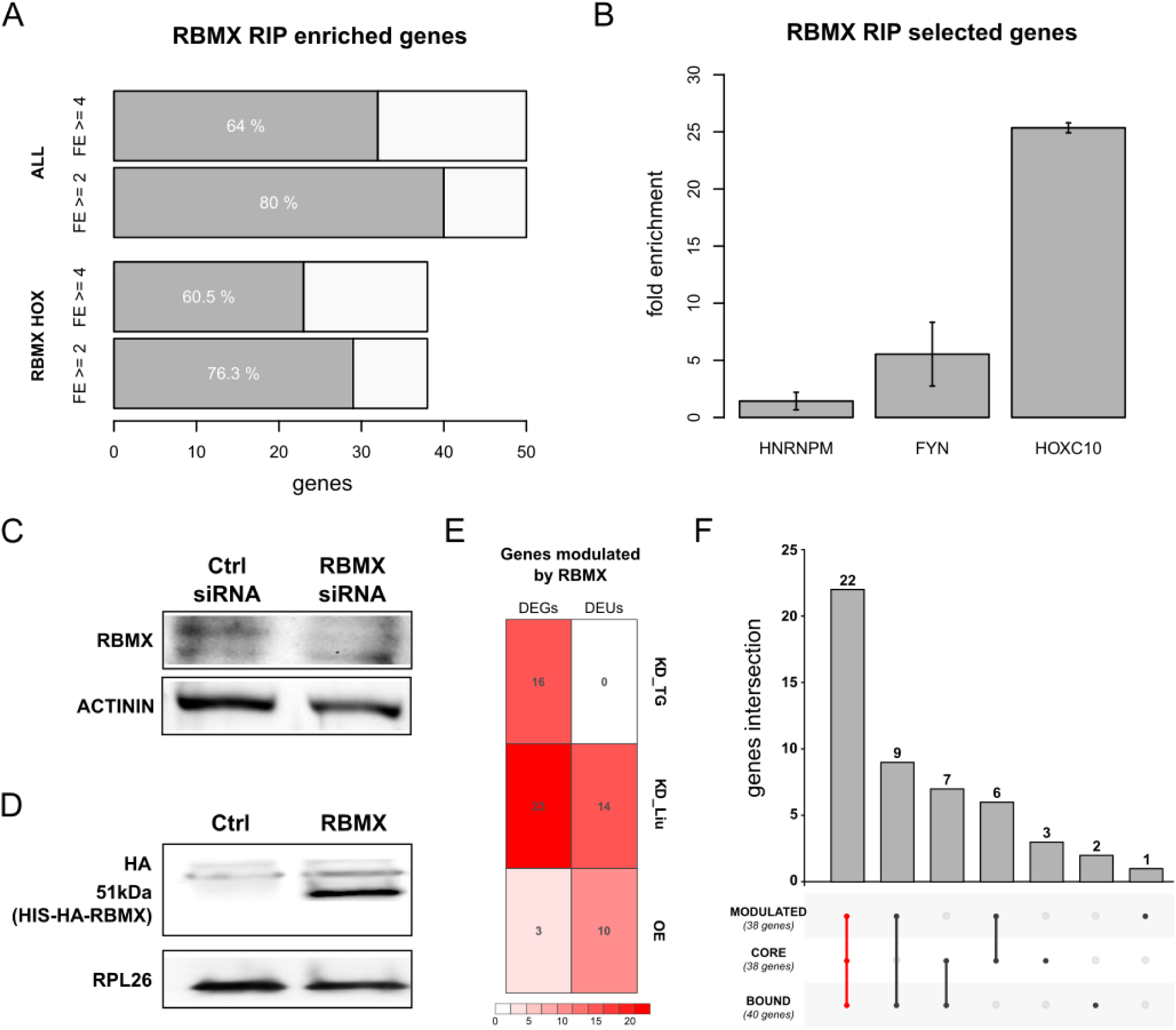
RBMX binds to homeotic genes mRNAs and controls their expression and splicing. **A)** shows the number of genes enriched in the RBMX RIP assay, at fold enrichment greater or equal than 2 (FE>=2) or 4 (FE>=4) in at least 2/3 replicates. Two sets of genes are considered, namely the core HOX genes set (RBMX HOX, 38 genes) and all tested genes (ALL, 50 genes). **B)** displays the fold enrichment in the RIP assay for the negative control (HNRNPM), the positive control (FYN), and one representative of the HOX core genes set (HOXC10). **C)** RBMX western blot in HEK293 cells treated with control and RBMX siRNA, with Actinin used as reference protein. **D)** HA-tag western blot in control and RBMX-overexpressing HEK293 cells, with RPL26 used as reference protein. **E)** shows the number of differentially expressed (DEGs) and differential exon usage (DEUs) genes in our RBMX knock-down targeted RNA-seq (KD_TG), the Liu RBMX knock-down dataset (KD_Liu, (Liu et al. 2017)) and our RBMX overexpression dataset (OE). **F)** displays all the intersections for the set of genes modulated by RBMX (MODULATED, defined as the union of DEGs and DEUs), the core set of homeotic genes (CORE), and genes bound by RBMX as per the RIP assay (BOUND). The intersection of all three is highlighted in red.

### RBMX controls homeotic genes mRNAs by post-transcriptional mechanisms

Given that RBMX binds to the mRNA of most homeotic genes containing a 5’HCE, we eventually sought to understand the impact this RBP has on their expression. To do so, we first used siRNAs to knock-down RBMX in HEK293 cells, reducing its protein level by 78% (**Figure 3C**, t-test p-value=0.00214). We thus performed a translatome profiling followed by targeted RNA-sequencing of total and polysomal fractions for the 50 genes which we previously tested by RNA immunoprecipitation. By this preliminary analysis, we identified 16 genes which were significantly upregulated at the polysomal level when silencing RBMX (log2 fold change >=1 and adjusted p-value <=0.1). Of these, 12 were part of the core homeotic genes and included members of all four HOX clusters, the DLX and POU families. However, replicates of samples at the total level were quite variable (average Spearman correlation=0.83). Furthermore, this assay does not allow to detect alternative splicing events, which can also be modulated by RBMX (Heinrich et al. 2009; Wang et al. 2011). So, to expand this analysis, we reanalyzed a recently published whole-transcriptome RNA-sequencing of HEK293 cells at the total level after RBMX knock-down (Liu et al. 2017). Eventually, we completed this dataset by overexpressing RBMX via a HIS-HA-tagged construct (**Figure 3D**) in the same cell type. These cells were then subjected to translatome profiling followed by whole-transcriptome RNA-sequencing of the total and polysomal fractions. Both datasets were analyzed to find differentially expressed genes (DEGs) and differential exon usage (DEU) events (**Figure 3E**). DEGs were mostly in the knock-down (23/50 genes, 6 up- and 7 down-regulated), with only 3/50 genes in overexpressed samples (two up- and one down-regulated, 2 DEGs at both the total and polysomal level, adjusted p-value <= 0.05). DEU events were more evenly distributed, with 10/50 genes affected in the overexpressed (all upregulated exons, 6 at both the total and polysomal level) and 14/50 in the knock-down (4 up-, 3 down-regulated and 7 with both up- and down-regulated exons, adjusted p-value <= 0.05). Eventually, we intersected all the datasets, also including the RIP data to identify a consistently controlled set of homeotic genes. As shown in **Figure 3F**, 38/50 genes (76%) are controlled by RBMX, via alternative splicing or differential expression at the total and polysomal level. Of these, 31 (62%) were also identified as bound by RBMX in the RIP assay. Of the 38 core homeotic genes, 28 are modulated (73%), 22 of which (58%) were also identified as bound by RBMX. Modulated genes in the core set include genes from all four HOX clusters, the DLX, NKX2 and POU family. Lists of modulated genes for each dataset can be found in **Supplementary Table 6** and **7**. Globally, this data indicates that RBMX extensively controls the fate of homeotic genes mRNAs through complementary regulatory mechanisms.

### RBMX is a translational regulator

RBMX controls its targets both by modulating their alternative splicing (Heinrich et al. 2009; Wang et al. 2011; Liu et al. 2017) and transcript abundance (Liu et al. 2017; Shin et al. 2008). The data we presented further confirm these findings. Furthermore, a few events (1 DEG and 4 DEUs) were observed at the polysomal level only, suggesting the possibility that RBMX may also act as a translational regulator. To verify this possibility, we thus checked whether RBMX is located in polysomes. We performed a polysomal profiling assay, followed by a fraction-by-fraction western blot against RBMX and RPL26, a component of the ribosome. As shown in **Figure 4A**, cytoplasmic RBMX is predominantly found in the polysomal fractions, with no apparent increase in abundance in heavy polysomes with respect to lighter ones.

**Figure 4.**
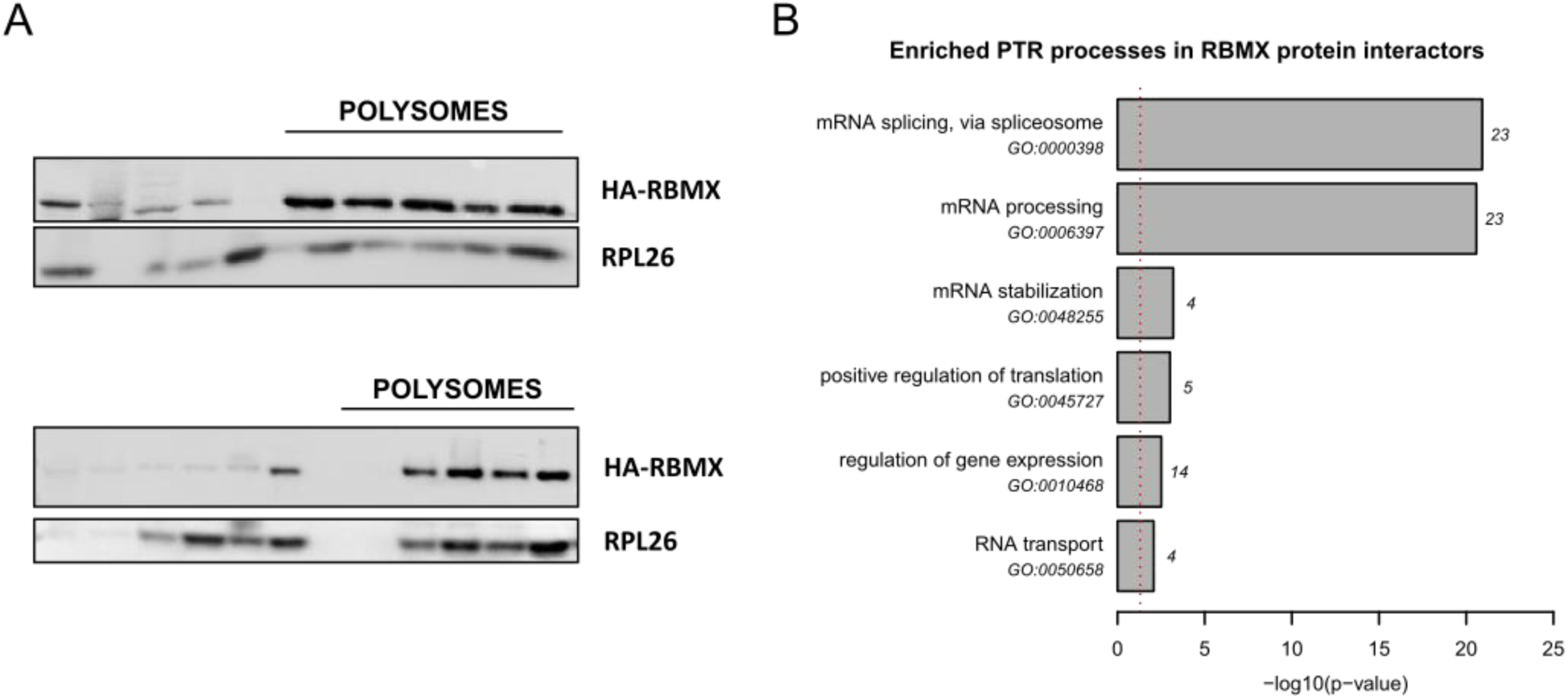
RBMX is a translational regulator. **A)** shows the distribution of RBMX on polysomes through a western blot of the fractions derived by polysomal profiling. The RPL26 ribosomal protein is used as positive control. **B)** displays post-transcriptional regulatory processes enriched in the set of RBMX protein-protein interactors. The enrichment p-value is shown on the x-axis as -log10(p-value). The number of RBMX interactors annotated to each process is instead shown next to the corresponding bar.

Eventually, we reasoned that observing RBMX associated with translation factors and ribosomal proteins would further suggest it to be a translational regulator. To verify this possibility, we thus analyzed known protein-protein interactions (PPIs) of RBMX. We collected experimentally determined PPIs from the STRING (Szklarczyk et al. 2017) and IntAct (Orchard et al. 2014) databases and performed a functional enrichment analysis of the resulting 83 RBMX interactors (full list of genes and enriched terms in **Supplementary Table 8** and **9**). As shown in **Figure 4B**, mRNA splicing and processing are enriched (23 genes, adjusted Fisher test p=1.21E-21 and 2.6E-21), along with mRNA stabilization (4 genes, adjusted Fisher test p=6.3E-04). RBMX interactors also include several translational regulators, represented by the “positive regulation of translation” process (5 genes, adjusted Fisher test p=9.7E-04). In particular, RBM3 interacts with RPL4 and is associated with polysomes through the 60S ribosomal subunit (Dresios et al. 2005), as is FXR2 (Siomi et al. 1996; Corbin et al. 1997). Eventually, CIRBP interacts with the EIF4G1 translation initiation factor (Yang et al. 2006), and KHDRBS1 associates with polysomes (Paronetto et al. 2006). Globally, these results show that RBMX can also act as a translational regulator.

### RBMX controls genes associated with XLID phenotypes

A frameshift deletion in the RBMX gene has been previously connected to the insurgence of an XLID syndrome associated with craniofacial dysmorphisms and intellectual disability, the Shashi syndrome (Shashi et al. 2015). We thus sought to check whether the homeotic genes we identified as modulated by RBMX could be involved with such phenotypes. To this end, we annotated the 22 homeotic genes in the core set which are bound and regulated by RBMX with three disease ontologies (see Methods). The results highlight several genes (HOXA2, NKX2-1, NKX2-5, POU2F1, and SATB2) associated with phenotypes compatible with this syndrome. These include mental retardation (SATB2), intellectual disability (NKX2-5), coarse facial features (NKX2-1 and NKX2-5), bulbous nose (SATB2), underdeveloped supraorbital ridges (NKX2-5), orofacial cleft (POU2F1, SATB2, HOXA2) and mixed hearing impairment (HOXA2). If considering all 31 modulated and bound genes, HOXB3, HOXB5, and SIX6, associated with obesity and microphthalmia, are also included. Eventually, we expanded this analysis to phenotypes commonly associated with XLID syndromes in general (Stevenson et al. 2013). Further genes (DLX5, HOXC4, HOXC5) are associated with compatible phenotypes such as cardiac malformations, synostosis, hypospadias, and renal anomalies. The list of all annotations can be found in **Supplementary Table 10**. Globally, the regulatory activity of RBMX could thus be strongly associated with this group of diseases.

## Discussion

In this work, we used a computational approach to extract phylogenetically hyper-conserved elements from the 5’UTR of human messenger RNAs (5’HCEs). We thus expanded the known catalog of regulatory mechanisms mediating the role of these regions in determining cell phenotypes. While much attention has been devoted to mapping functional elements in the 3’UTR, its 5’ counterpart is still relatively uncharacterized. Our approach focused on extracting the most highly conserved regions, under the assumption that these would be evolutionary stable PTR sites of utmost importance.

The 5248 5’HCEs we identified are short regions occurring in around 10% of protein-coding genes, most often localized in proximity to one another. Given their prevalently clustered nature, 5’HCEs could represent loci of cooperation and competition between post-transcriptional regulatory factors. Through their interplay, these factors would ultimately determine the translation of the containing mRNAs. As 5’HCEs do not systematically overlap with uORFs or IRESes, binding sites for the about 1500 human RNA-binding protein (RBP) genes are the likely orchestrators of such behaviors. Understanding how these mechanisms work and impact cell physiology and pathology will thus require further efforts towards the systematic mapping of RBP binding sites.

Among genes whose mRNA contain a 5’HCE, we identified a broad set of homeotic genes including members of all HOX clusters and several other related families. Homeotic genes are the prototypical class of conserved genes in metazoa, responsible for the development of the body plan, organs, and the nervous system (Holland 2013; Philippidou and Dasen 2013; Mallo et al. 2010). In that respect, they represent the ideal result of our algorithm, benchmarking its ability to identify truly conserved regions. While their transcriptional regulation is well characterized, we still lack a clear picture of how homeotic genes are controlled at the post-transcriptional level. In particular, only a few studies have explored the role of 5’UTRs in their regulation (Regadas et al. 2013; Nie et al. 2015; Xue et al. 2015). Identifying this set of 5’HCEs thus allowed us to improve our understanding of how PTR controls development through the 5’UTRs.

By applying de novo motif search combined with known RBP motifs matching we identified RBMX as a candidate post-transcriptional regulator of homeotic genes. Phenotypic studies have shown this protein, which is itself highly conserved, to affect neuromuscular development in *X. laevis* (Dichmann et al. 2008) *and D. rerio* (Tsend-Ayush et al. 2005). This evidence pointed to a possible role for RBMX as a controller of mRNAs of homeotic genes. We confirmed this hypothesis by a combination of RNA immunoprecipitation, knock-down, overexpression experiments, and sucrose-based cytoplasmic fractionation. This assays allowed us to probe multiple aspects of post-transcriptional regulation, including alternative splicing, mRNA stabilization, and translation. From this data, we found a possible new role for RBMX as a translational regulator. This finding establishes RBMX as a versatile controller of the mRNA, able to impact its lifecycle from alternative splicing to protein production. This flexibility may have evolved to allow better control of processes requiring particularly fine tuning such as body plan establishment and neural development.

We have shown that RBMX regulates the mRNAs of many homeotic genes. However, other RBPs may also contribute to the post-transcriptional regulation of homeotic genes, possibly by cooperating or competing with RBMX. Further studies will thus be needed to complete this regulatory network.

Eventually, we explored the functional implications stemming from the role of RBMX. Which phenotypes does it drive? How would these be affected if its action was perturbed by genetic alterations typical of many diseases? An example is the Shashi syndrome (Shashi et al. 2015): a frameshift deletion in RBMX is associated with the onset of this XLID disease. Indeed, several of the homeotic targets of RBMX we identified are responsible for compatible phenotypes, such as mental retardation and cardiac malformations. Abnormal RBMX expression and function could thus lead to the insurgence of such syndromes through downstream altered expression of homeotic genes. Tuning the post-transcriptional networks RBMX controls could offer an important therapeutic opportunity for this class of diseases and, possibly, other developmental pathologies. Further studies are thus warranted to complete the characterization of this RBP as a master regulator of development.

## Materials and Methods

### 5’HCE identification and characterization

Human 5’UTRs, the related 44-vertebrates alignment and sequence conservation scores (SCS) were downloaded from UCSC for the hg18 genome assembly (Tyner et al. 2017). Branch length score (BLS) was computed for each UTR as described in (Dassi et al. 2013), and composed at equal weight with SCS to derive the hyper-conservation score (HCS). Then, a sliding-window algorithm, starting with fully conserved 5-nucleotides seeds (HCS = 1.0) and expanding them upstream and downstream until a minimum score threshold of 0.85 is reached, was applied to the 5’UTRs, as described in (Dassi et al. 2013), to derive hyper-conserved elements (HCE).HCE properties were then computed by custom Python scripts and plotted with R. Functional enrichment in the set of genes containing HCEs in their 5’UTR was computed by DAVID (Huang et al. 2007) using Gene Ontology (BP, MF, and CC parts) and protein domain ontologies (INTERPRO, PFAM, and SMART).

### Motif analysis

De novo motif search was performed on the sequences of 5’HCEs from homeotic genes using DynaMIT (Dassi and Quattrone 2016), configured to use two search tools: Weeder (Pavesi et al. 2004) for sequence motifs (lengths 6, 8, 10 and 12 nts with 1, 2, 3 and 4 mismatches allowed resp.; at least 25% of the sequences containing the motif) and RNAforester (Höchsmann et al. 2003) for secondary structure motifs (multiple alignment and local search modes). The selected motif integration strategy was “co-occurrence”, which computes the co-occurrence score between motifs pairs as the Jaccard similarity of the mutual presence of both motifs on each sequence. This measure thus allows finding motifs which co-regulate the same sequences set. The best motif from DynaMIT result was selected, and only positions with at least ten supporting sequences were kept. This resulted in trimming the 5’ and 3’ ends of the integrated motif. These trimmed ends represented lowly-supported, more “peripheral” individual motifs with respect to the core part consistently shared by multiple motifs.

The PWMs for a set of 193 RBPs were obtained from the CISBP-RNA database (Ray et al. 2013). Pearson correlation between the CISBP-RNA PWMs and the motif identified by DynaMIT were computed by the TFBSTools R package (Tan and Lenhard 2016). The RBMX PWM obtained from CISBP-RNA was matched against the sequences of 5’HCEs from homeotic genes using a custom Python script and the Biopython library (Cock et al. 2009). Only PWMs at least four nucleotides long were used, and only matches with a score greater than 70% were considered.

### Cell culture and transfection

HEK293 cells were cultured in DMEM with 10% FBS, 100 U/ml penicillin-streptomycin and 0.01 mM l-glutamine (Gibco, Waltham, MA). Cultures were maintained at 37°C in a 5% CO2 incubator.

### RBMX knock-down and overexpression

We performed RBMX knock-down as described in (Matsunaga et al. 2012). Briefly, we used RBMX siRNA-1 (5’-UCAAGAGGAUAUAGCGAUATT-3’) and RBMX siRNA-2 (5’-CGGAUAUGGUGGAAGUCGAUU-3’) for RBMX knock-down, and negative control siRNA S5C-060 (Cosmo Bio, Tokyo). 1.5×10^6^ HEK293 cells were seeded into two 10-cm Petri dishes and transfected with a mixture of both siRNA at 25nM using Lipofectamine 2000.

Full-length RBMX was amplified by PCR using HeLa cells cDNA and the following primers: Fw: 5’ GAGGCGATCGCCGTTGAAGCAGATCGCCCAGGAA 3’ and Rv: 5’ GCGACGCGTCTAGTATCTGCTTCTGCCTCCC 3’. The amplified fragment was digested with the SgfI and MluI restriction enzymes and cloned into the pCMV6-AN-His-HA plasmid (PS100017, OriGene, Rockville, MD) to obtain the pCMV6-HIS-HA-RBMX vector, expressing the gene fused with an amino-terminal polyhistidine (His) tag and a hemagglutinin (HA) epitope. The construct was confirmed by sequencing. 1.5×10^6^ HEK293 cells were seeded into two 10-cm Petri dishes and transiently transfected using Lipofectamine 2000 (Invitrogen, Waltham, MA) with 2µg of pCMV6-HIS-HA-RBMX or the mock empty vector as control. Total and polysomal RNA extractions were performed 48h post-transfection. All the experiments were run at least in biological triplicate.

### RNA immunoprecipitation

Ribonucleoprotein immunoprecipitation (RIP) was performed in three biological replicates using lysates of human HEK293 cells transfected with pCMV6-HIS-HA-RBMX or with the mock empty vector. Cell extracts were resuspended in NT2 buffer (50 mM Tris–HCl pH 7.4, 150 mM NaCl, 1 mM MgCl2, 0.05% NP-40 supplemented with fresh 200 U RNase Out, 20 mM EDTA and a protease inhibitor cocktail), chilled at 4°C. Anti-HA magnetic beads (Pierce, Waltham, MA, USA) were saturated in the NT2 buffer (adding 5% BSA for 1h at 4°C), then added to lysates. The immunoprecipitation was performed overnight at 4°C in gentle rotation condition. Eventually, the immunoprecipitate was washed four times with NT2 and resuspended in the same buffer. RNA extraction was performed from 10% of the volume of both the input and the immunoprecipitate samples, using TRIzol (Invitrogen, Carlsbad, CA, USA). Sequencing libraries were then prepared following the manufacturer’s instructions as described below. FYN mRNA was used as the positive control and HNRNPM as the negative control (Heinrich et al. 2009).

### Polysomal fractionation and RNA extraction

Cells were incubated for 4min with 10 µg/ml cycloheximide at 37°C to block translational elongation. Cells were washed with PBS + 10 µg/ml cycloheximide, scraped on the plate with 300 µl lysis buffer (10 mM NaCl, 10 mM MgCl2, 10 mM Tris-HCl, pH 7.5, 1% Triton X-100, 1% sodium deoxycholate, 0.2 U/µl RNase inhibitor [Fermentas Burlington, CA], 10 µg/ml cycloheximide, 5 U/mL DNase I [New England Biolabs, Hitchin, UK] and 1 mM DTT) and transferred to a tube. Nuclei and cellular debris were removed by centrifugation for 5min at 13,000g at 4°C. The lysate was layered on a linear sucrose gradient (15-50% sucrose (w/v), in 30 mM Tris–HCl at pH 7.5, 100 mM NaCl, 10 mM MgCl2) and centrifuged in an SW41Ti rotor (Beckman Coulter, Indianapolis, IN) at 4°C for 100min at 180,000g. Ultracentrifugation separates polysomes by the sedimentation coefficient of macromolecules: gradients are then fractionated and mRNAs in active translation, corresponding to polysome-containing fractions, separated from untranslated mRNAs. Fractions of 1 mL volume were collected with continuous monitoring absorbance at 254 nm. Total RNA was obtained by pooling together 20% of each fraction. To extract RNA, polysomal and total fractions were treated with 0.1 mg/ml proteinase K (Euroclone, Italy) for 2h at 37°C. After phenol-chloroform extraction and isopropanol precipitation, RNA was resuspended in 30 µl of RNase-free water. RNA integrity was assessed by an Agilent Bioanalyzer and RNA quantified by a Qubit (Life Technologies, Waltham, MA).

### Protein extraction and Western blots

10% of each fraction collected from sucrose gradient fractionation was pooled (for knock-down and overexpression validation) or processed separately (for the RBMX polysomes distribution assay) to extract proteins using TCA/acetone precipitation. Proteins were resolved on 15% SDS-PAGE, transferred to nitrocellulose membrane and immunoblotted with RBMX (Abcam, Cambridge, UK), HA (Bethyl Laboratories, Montgomery, TX) and RPL26 antibodies (Abcam, Cambridge, UK). Blots were processed by an ECL Prime detection kit (Amersham Biosciences).

### RNA-seq

For the RNA immunoprecipitation and the RBMX knock-down samples, after total and polysomal RNA extraction, 500ng RNA of each sample were used to prepare libraries according to the manufacturer’s protocol, using a TruSeq Targeted RNA Custom Panel Kit (Illumina, San Diego, CA). Sequencing was performed with a 50-cycle MiSeq Reagent Kit v2 (Illumina, San Diego, CA) on a MiSeq machine. For the overexpression samples, after total and polysomal RNA extraction, 500ng RNA of each sample were used to prepare libraries according to the manufacturer’s protocol, using the TruSeq RNA Sample Prep Kit (Illumina, San Diego, CA). Sequencing was performed on six lanes at 2×100bp on a HiSeq 2500 machine. Sequencing was performed at least in triplicate for each sample.

### RNA-seq data analysis

Reads were pre-processed with trimmomatic (Bolger et al.), trimming bases having quality lower than Q30 and removing sequencing adapters. Remaining reads were aligned to the human genome hg38 assembly with bowtie2 (Langmead and Salzberg 2012) for the targeted sequencing, and with STAR for the other datasets (Dobin et al. 2013). Read counts for each gene determined using the Gencode v27 annotation (Harrow et al. 2012), and normalized by the sequencing depth of the libraries. Fold enrichment for RNA immunoprecipitation were computed as the ratio of enrichment of (RIP vs. input samples for the RBMX IP) over (RIP vs. input samples for the HA IP), for each replicate. For the targeted RNA-seq of RBMX knock-down samples, samples with less than 500 mapped reads (less than 10 mapped reads per targeted gene) were discarded, and differences in expression were computed by a Wilcoxon test, as this type of library (probing only 50 genes) does not fit the assumptions of commonly used differential expression determination methods. For overexpression and the published RBMX knock-down we obtained from (Liu et al. 2017), differential expression was computed with DESeq2 (Love et al. 2014), while differential exon usage was obtained by DEXseq (Anders et al. 2012) (adjusted p-value <=0.05 for both analyses). Functional annotation was performed by Enrichr (Chen et al.) on Gene Ontology or disease ontologies (OMIM disease, Jensen disease, and Human Phenotype Ontology) annotations.

## Data availability

Datasets can be found in GEO with ID GSE118383 (RBMX RIP targeted RNA-seq and RBMX knock-down targeted RNA-seq) and GSE68990 (RBMX overexpression RNA-seq).

## Supplemental figures

**Supplementary Figure 1:**
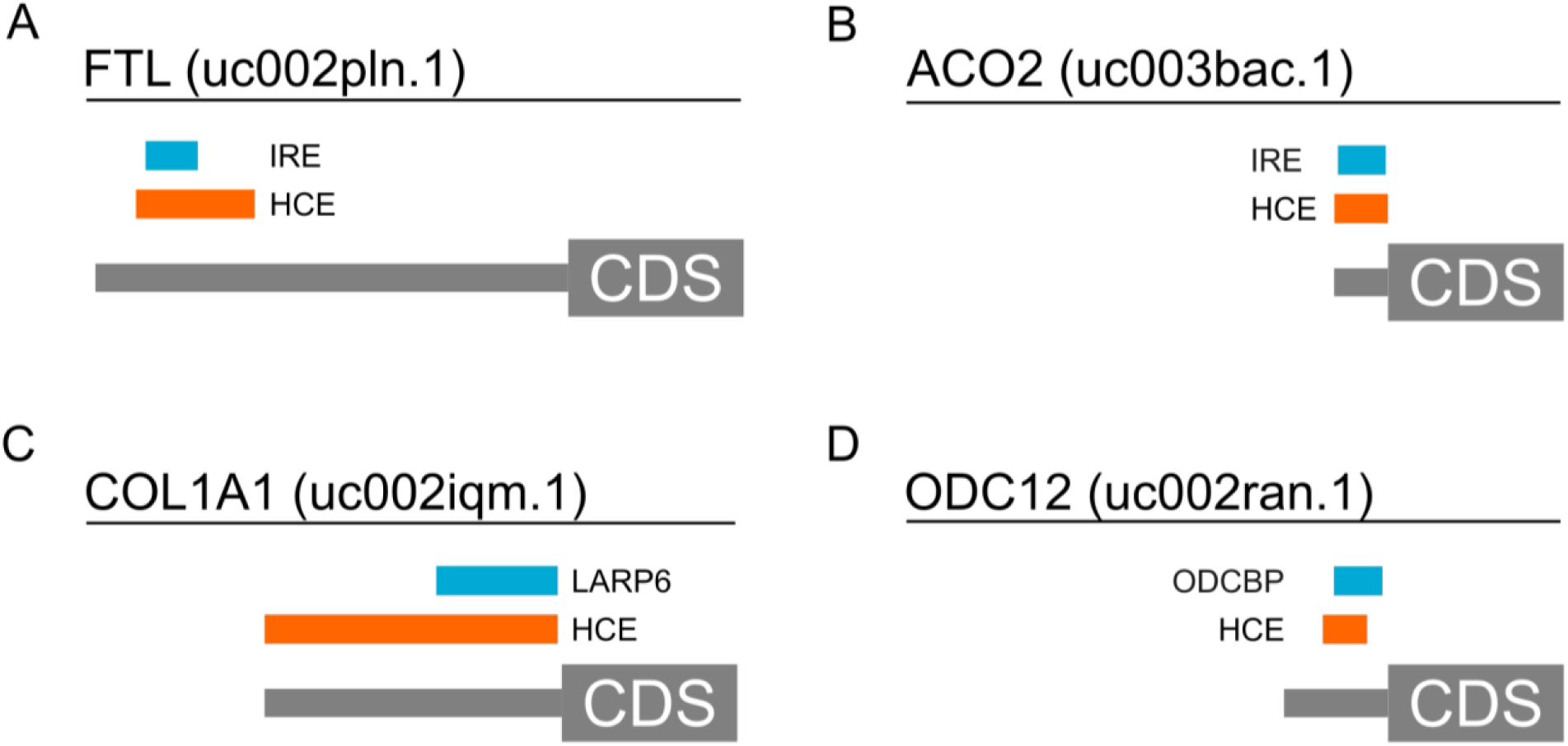
5’HCEs contain conserved binding sites and cis-elements. The figure displays four example of functional regions retrieved by the HCE identification algorithm in 5’UTRs. **A)** a known, conserved iron response element (IRE) in the 5’UTR of ferritin (FTL) is fully contained in an HCE. **B)** a known, conserved iron response element (IRE) in the 5’UTR of aconitase 2 (ACO2) is fully contained in an HCE. **C)** a conserved binding site for LARP6 in the 5’UTR of collagen alpha type 1 (COL1A1) is fully contained into an HCE. **D)** a conserved binding site for ODCBP in the 5’UTR of ornithine decarboxylase 1 (ODC1) is partially overlapping with an HCE.

## Acknowledgements

We thank Veronica De Sanctis and Roberto Bertorelli (NGS Facility, Centre for Integrative Biology and LaBSSAH, University of Trento) for performing NGS sequencing.

